## Supplementary Information for "MultiCens: Multilayer network centrality measures to uncover molecular mediators of tissue-tissue communication"

**for the manuscript "MultiCens: Multilayer network centrality measures to uncover molecular mediators of tissue-tissue communication"**

#### 1 Supplementary Results

##### 1.1 Literature support for our hormone-gene predictions – additional information

We discuss here the predictions for the growth hormone Somatotropin to supplement similar discussions in the Results section in the main text. To add to examples of novel predictions that are not in ground truth HGv1 and also have poor PubMed literature support scores, we discuss *S100A8* for Somatotropin – there is not substantial literature support for this prediction, but there are studies that show downregulation of *S100A8* on exogenous administration of the growth hormone [1]. *EGFR* (Epidermal growth factor receptor) playing key roles in development, cellular proliferation, and cancer, can be modulated by growth hormone [2,3]. Increased level of *FFAR4* (free fatty acid receptor 4) reduces ghrelin secretion, further stimulating hunger [4]. This context-based dependency is well captured in the cosine-based similarity in the embedding space, but the gene has low or no direct co-occurrence with hormone-related terms (see Fig. 4b of main text).

##### 1.2 Literature support for our hormone-lncRNA predictions – additional information

We discuss here the long non-coding RNA (lncRNA) predictions for different hormones to supplement similar discussions in the Results section in the main text. Since Suppl Table S3 lists all supporting references for each hormone-lncRNA prediction, we do not cite all these references in the text below, similar to what we do in the main text for better readability.

We first discuss lncRNA predictions related to insulin. Our centrality-based ranking of insulin-relevant pancreas lncRNAs revealed *HOXA-AS2*, which has links to diabetes through another gene *TIMP3* [5]. Further, interplay between lncRNA and insulin pathway related genes are indicated in pathogenesis of different diseases [6]. This phenomenon is supported by our lncRNA prediction in pancreas, where most of them promote tumorigenesis and metastasis (*LINC00672* promotes endometrial cancer chemosensitivity, *HOXA-AS2* promotes cellular processes aiding non-small cell lung cancer or pancreatic cancer, *PRR34-AS1* is highly expressed in hepatocellular carcinoma, and *LINC00294* induced by glucose-regulated protein 78, *GRP78*, aids in advancement of cervical cancer). Similarly, lncRNAs predicted to be present in skeletal muscle are also involved in tumorigenesis (*ZEB1-AS1* promotes pancreatic cancer progression, *TNK2-AS1*/miR-125a-5p fosters the progression of gastric cancer, whereas *PWAR6* and *PRRT3-AS1* act as tumour suppressors in glioma and prostate cancer respectively).

For the growth hormone Somatotropin, many predicted lncRNAs have connections to cancer cell growth as described next. While loss of *LINC01132* attenuates ovarian tumour growth, silencing lncRNA-*UCA1* leads to repression of pituitary cancer cell growth and prolactin (PRL) secretion and *LINC01473* expression is negatively correlated with serum interleukin-2 and tumor necrosis factor $\alpha$ levels in multiple myeloma. Similarly, *PTPRD-AS1* is inversely correlated with the overall survival in patients with ovarian cancer. Further, *PTPRD* (receptor protein tyrosine phosphatase delta) hinders the growth of glioblastoma multiforme and other tumor cells, and also human astrocytes in dearth of *PTPRD* exhibits growth escalation. This probably indicates that *PTPRD-AS1*, by controlling PTPRD expression level, helps in glial cell growth.

LncRNAs found with high progesterone-specific query set centrality are involved in several cancers, including colon adenocarcinoma (*TAF1A-AS1*), prostate cancer (*PCAT19*), ovarian cancer (*HHIP-AS1*), breast cancer (*LINC00641*, *MIR210HG*, *HAGLR*), endometrial cancer (*MIR210HG*, *LINC01016*), and many others.

##### 1.3 MultiCens analysis of AD vs. CTL networks using a different query set

We showed in main text how the change in the four-brain-region gene networks between Alzheimer's disease (AD) vs. Control (CTL) groups can be investigated by applying MultiCens with synaptic signaling genes (also referred to as synaptic genes or SSG) as the query set. When we changed the synaptic signaling gene set (SSG, 134 genes) to Plaque-induced gene set (PIG, 57 genes) for the query set, MultiCens centralities of SSG vs. PIG were highly but not perfectly correlated (see Suppl Fig. S4). This resulted in top-ranking genes and pathway enrichments for PIG, some of which are similar between PIG and SSG as discussed first below, and others that are different. Brain region specific similarities and differences were also noted. For instance, for PIG, we see that HSP90 chaperone cycle for steroid hormone receptors (SHR) pathway is again enriched in AD group in all 3 brain regions as for SSG. Similarly biological process "protein folding" is found to be enriched in both PIG and SSG set in AD group. Moreover, *JMJD6*, *SLC5A3*, *CIRBP*, and *AHSA1* are also among the top ten genes in AD group (as for SSG; see Results in main text for SSG top-ranking genes). Pathway related to extracellular matrix (ECM) organization (R-HSA-1474244) is also enriched for correlation to PIG genes in AD brain regions. The ECM is known to contribute to both A$\beta$ plaques' formation and degradation [7]. In case of control group, pathway related to immune system and biological process concerning cytokine production are positively enriched. A$\beta$ is a known constituent of the innate immune system and regarded as an "early responder cytokine" [8]. The change in gene ranking and pathway enrichments for top ranks for PIG relative to SSG is highlighted in Suppl Fig. S5a and S5b respectively (compare with Fig. 5 of main text). For instance, biological processes like "regulation of hemopoiesis" and pathways like "Serotonin Neurotransmitter Release Cycle",

"ER to Golgi Anterograde Transport" and "Interleukin-4 and Interleukin-13 signaling" were found in PIG-based, but not SSG-based, enrichment analysis. On the other hand, pathways like "Voltage gated Potassium channels", "Cell-cell junction organization" and "The role of GTSE1 in G2/M progression after G2 checkpoint" and biological processes like "axon development" were prominently enriched for SSG gene set.

#### 2 Supplementary Methods

**Definition 1.** *Local centrality vector is defined as the following iterative equation*

$$l = pAl + \frac{(1-p)}{n}\vec{1} \tag{1}$$

**Definition 2.** *For a given local centrality l, global centrality vector in a multi-layer network can be defined by the following iterative equation*

$$g = p\Big[(A+C)g + Cl\Big] + \frac{(1-p)}{N}\vec{1} \tag{2}$$

##### 2.1 Convergence and Decomposability of the proposed centrality measures

In this section, we first discuss the convergence of the proposed centrality measures. Proof of convergence of local and global centrality, and how they constitute to versatility is given in the main text as Theorem one and Theorem 2.

**Definition 3.** *For a given l, layer-specific centrality vector in a multi-layer network can be defined by the following iterative equation*

$$g_{layer(i)} = p\Big[(A+C)g_{layer(i)} + C^{[i]}l\Big] + \frac{(1-p)}{N}\vec{1} \tag{3}$$

*where $C^{[i]}$ represents the matrix $C$ with all but ith column-block entries set to* 0*.*

**Theorem 3** *For $0 \le p < 1$, $g_{layer(i)}$ defined by Equation 3 always converges.*

*Proof.* From equation 3,

$$\begin{aligned}
g_{layer(i)} &= p\Big[(A+C)g_{layer(i)} + C^{[i]}l\Big] + \frac{(1-p)}{N}\vec{1} \\
&= p\left[(A+C)\Big(p\big[(A+C)g_{layer(i)} + C^{[i]}l\big] + \frac{(1-p)}{N}\vec{1}\Big) + C^{[i]}l\right] + \frac{(1-p)}{N}\vec{1} \\
&= p\left[p(A+C)^2 g_{layer(i)} + p(A+C)C^{[i]}l + (A+C)\frac{(1-p)}{N}\vec{1} + C^{[i]}l\right] + \frac{(1-p)}{N}\vec{1} \\
&\ \ \vdots \\
&= p^k(A+C)^k g_{layer(i)} + p\sum_k p^k(A+C)^k C^{[i]}l + \sum_k p^k(A+C)^k\frac{(1-p)}{N}\vec{1} \\
&\quad + \frac{(1-p)}{N}\vec{1}
\end{aligned}$$

**Theorem 4** *Global centrality, as defined in the main text, can be decomposed for each layer being a specific target layer.*

$$\sum_{i=1}^{L} g_{layer(i)} = g \tag{4}$$

*Proof.*

$$\sum_{i=1}^{L} g_{layer(i)} = p\Big[\big(A+C\big)\sum_{i=1}^{L} g_{layer(i)} + LCl\Big] + \frac{L(1-p)}{N}\vec{1}$$

$$\tilde{g} = p\Big[\big(A+C\big)\tilde{g}\Big] + L\Big[pCl + \frac{(1-p)}{N}\vec{1}\Big]$$

$$\tilde{g} = Lg$$

Since $g$ is a centrality vector, $L$ being a constant can be ignored. So the formulation can be written as,

$$\sum_{i=1}^{L} g_{layer(i)} = g$$

This completes the proof.

**Definition 4.** *For a given set of query nodes $set(k)$, the local-set centrality in a multilayer network can be defined by the following equation.*

$$l^{set(k)} = pAl^{set(k)} + \frac{(1-p)}{n}\vec{1^k} \tag{5}$$

*Where $\vec{1^k}$ represents the vector of $1'$s at indices corresponding to the node-set $k$ and 0 otherwise.*

Let $\{k_1, k_2, \ldots, k_K\}$ denote the collectively exhaustive subsets of nodes in a layer, and without loss of generality, $k$ denotes any subset from this set. Now we show that $l^{set(k)}$ converges, and for a given layer, $l^{set(k)}$ adds up to $l$.

**Theorem 5** *For $0 \leq p < 1$, $l^{set(k)}$ defined by Equation 5 always converges.*

*Proof.* From equation 5,

$$l^{set(k)} = pAl^{set(k)} + \frac{(1-p)}{n}\vec{1^k}$$

$$l^{set(k)} = pA\Big(pAl^{set(k)} + \frac{(1-p)}{n}\vec{1^k}\Big) + \frac{(1-p)}{n}\vec{1^k}$$

$$l^{set(k)} = p^2A^2l^{set(k)} + pA\frac{(1-p)}{n}\vec{1^k} + \frac{(1-p)}{n}\vec{1^k}$$

$$\vdots$$

$$l^{set(k)} = p^jA^jl^{set(j)} + \sum_{i=0}^{j-1} p^iA^i\frac{(1-p)}{n}\vec{1^k}, \text{ where } j \to \infty$$

The right side of the equation is similar to the original PageRank centrality which is known to converge for $0 \leq p < 1$.

**Theorem 6** *Local-set centrality defined by equation 5 can be added for each set $k$ to obtain the local centrality $l$.*

$$\sum_{k=1}^{K} l^{set(k)} = l \tag{6}$$

*Proof.*

$$\sum_{k=1}^{K} l^{set(k)} = pA \sum_{k=1}^{K} l^{set(k)} + \frac{(1-p)}{n} \sum_{k=1}^{K} 1^{\vec{k}}$$

$$\tilde{l} = pA\tilde{(l)} + \frac{(1-p)}{n} 1^{\vec{k}}$$

This equation is the same as the iterative equation defined for computing local centrality. This completes the proof.

We use this *local-set* centrality to define *query-set* centrality as follows.

**Definition 5.** *For a given set of query genes $set(k)$ in a layer $i$, the query-set centrality in a multilayer network can be defined by the following equation.*

$$g^{set(k)}_{layer(i)} = p\Big[(A+C)g^{set(k)}_{layer(i)} + C^{[i]} l^{set(k)}\Big] + \frac{(1-p)}{N} 1^{\vec{k}} \tag{7}$$

The *query-set centrality* is defined in order to capture the effect of nodes on a query set of genes in a specific target layer. Now we discuss the convergence of this centrality equation.

**Theorem 7** *For $0 \leq p < 1$, $g^{set(k)}_{layer(i)}$ defined by Equation 7 always converges.*

*Proof.* From equation 7,

$$\begin{aligned}
g^{set(k)}_{layer(i)} &= p\Big[(A+C)g^{set(k)}_{layer(i)} + C^{[i]} l^{set(k)}\Big] + \frac{(1-p)}{N} 1^{\vec{k}} \\
&= p\left[(A+C)\left(p\big[(A+C)g^{set(k)}_{layer(i)} + C[i] l^{set(k)}\big] + \frac{(1-p)}{N} 1^{\vec{k}}\right) + C^{[i]} l^{set(k)}\right] \\
&\quad + \frac{(1-p)}{N} 1^{\vec{k}} \\
&= p\left[p(A+C)^2 g^{set(k)}_{layer(i)} + p(A+C)C^{[i]} l^{set(k)} + (A+C)\frac{(1-p)}{N} 1^{\vec{k}} + C^{[i]} l^{set(k)}\right] \\
&\quad + \frac{(1-p)}{N} 1^{\vec{k}} \\
&\vdots \\
&= p^d (A+C)^d g^{set(k)}_{layer(i)} + p\sum_d p^d (A+C)^d C^{[i]} l^{set(k)} + \sum_d p^d (A+C)^d \frac{(1-p)}{N} 1^{\vec{k}} \\
&\quad + \frac{(1-p)}{N} 1^{\vec{k}}
\end{aligned}$$

Following the discussion from Theorem 3, it can be shown that the right-hand side of the equation results in multiple geometric series, and all of them converge as the number of iterations increases. This completes the proof.

We now discuss the decomposability of the proposed centrality measures. As shown in Fig. 1 of the main text, our proposed centrality measures follow a hierarchical organization (tree structure) and centrality measures defined at each level add up to the parent node.

Similar to the decomposition at the level of *local* and *global* centralities, we can show that the *layer-specific* centralities of all layers add up to the *global centrality* equation.

**Theorem 8** *Layer-specific centrality defined by equation 3 can be decomposed into query-set centrality defined over collectively exhaustive subsets of nodes.*

$$\sum_k g_{layer(i)}^{set(k)} = g_{layer(i)} \tag{8}$$

*Proof.*

$$\sum_k g_{layer(i)}^{set(k)} = p\Big[(A+C)\sum_k g_{layer(i)}^{set(k)} + C^{[i]}\sum_k l^{set(k)}\Big] + \frac{(1-p)}{N}\sum_k \vec{1^k}$$

$$\tilde{g}_{layer(i)}^{set(k)} = p\Big[(A+C)\tilde{g}_{layer(i)}^{set(k)} + C^{[i]}\sum_k l^{set(k)}\Big] + \frac{(1-p)}{N}\sum_k \vec{1^k}$$

By using theorem 6

$$\tilde{g}_{layer(i)}^{set(k)} = p\Big[(A+C)\tilde{g}_{layer(i)}^{set(k)} + C^{[i]} l\Big] + \frac{(1-p)}{N}\vec{1}$$

The right side of the equation is the same as equation 3. This completes the proof.

##### 2.2 Hyperparameters and method complexity

Following the conventions of PageRank centrality algorithm, we tested $p$ in the range of $[0.7, 0.95]$. The higher values of $p$ tend to assign higher centrality scores to genes that are part of communities as compared to smaller values of $p$. In all our experiments, we report results at $p = 0.9$. However, there is very small deviation of rankings between $p = 0.85$ (typical value of $p$ used for web-based networks) and $p = 0.9$.

In the proposed method, the following two steps are involved:

1. Construction of multilayer network
2. Centrality score computation

The first step of network construction can be performed using multiple ways. In this project, we opted for correlation-based methods; hence we calculate correlation for all possible gene-gene pairs. Assuming a multilayer network with $L$ layers and $n$ genes per layer, we need to compute the correlation between $(L \times n)^2$ pairs. Each correlation can be computed with $O(k)$ time complexity, where $k$ is the number of samples. So the total runtime complexity of network construction is $O(k(L \times n)^2)$. In this project, we worked predominantly with on two-layered networks with around 15k genes per layer. The number of samples can be in few hundreds depending upon the intersection of sample IDs between the tissues. In our experiments, it takes roughly three hours to generate a two-layered multilayer network. The network can be stored in an adjacency matrix or list format. We store these networks in an adjacency matrix format of size $(L \times n)^2$. In practice, the matrix file can take up to a few GBs of memory.

The second step, which is the major contribution of work - the centrality computation, uses iterative equations to find the scores. Each iteration takes $O(nL \times nL)$ computations. The number of iterations depends upon various factors such as the diameter of the graph, modularity of the graph, etc. In practice, the method converges under 50 iterations incurring a total time of around thirty minutes. This step can be made efficient by distributing the code to multiple machines similar to distributed PageRank [9].

#### 3 Supplementary Tables

| **S. No.** | **Hormone** | **Gene-set (size)** | **AUC (coexp)** | **AUC (coexp + SNAP)** |
|---|---|---|---|---|
| 1 | Adrenaline | Target (24) | 0.469 | 0.494 |
| 2 | Aldosterone | Source (11) | 0.458 | 0.458 |
| 3 | Angiotensin | Target (15) | 0.507 | 0.557 |
| 4 | Cortisol | Source (12) | 0.533 | 0.542 |
| 5 | Estradiol | Target (89) | 0.493 | 0.512 |
| 6 | Glucagon | Target (19) | 0.552 | 0.574 |
| 7 | Insulin | Source (156) | 0.668 | 0.677 |
| 8 | Insulin | Target (215) | 0.664 | 0.685 |
| 9 | Norepinephrine | Source (16) | 0.473 | 0.475 |
| 10 | Norepinephrine | Target (14) | 0.430 | 0.471 |
| 11 | Progesterone | Source (13) | 0.740 | 0.733 |
| 12 | Progesterone | Target (35) | 0.598 | 0.645 |
| 13 | Somatotropin | Source (10) | 0.679 | 0.712 |
| 14 | Somatotropin | Target (22) | 0.564 | 0.671 |
| 15 | Thyroxin | Source (12) | 0.482 | 0.498 |
| 16 | Vitamin-D | Target (41) | 0.570 | 0.578 |

Table S1: Area under recall-at-k curve (AUC) for the ranking obtained using MultiCens query-set centralities, which were computed in the hormone-related human multilayer networks' application. For comparison, AUC for a random ranking of all genes is 0.5. We evaluated only hormones with at least 10 genes on the source or target tissue side, so that these gene sets to be retrieved are sufficiently large to yield a reliable estimate of AUC (see Suppl Fig. S2 and Fig. 3a of main text for recall-at-k curves; see also Fig. 3b (Coexpression+SNAP based results) of main text for more information about this application/evaluation, and a visualization of this table).

| (a) Insulin: Pancreas (predictions of insulin-producing genes) | |
|---|---|
| Top genes | Gene names |
| *SYBU* | syntabulin |
| *LRP1* | LDL receptor related protein 1 |
| *CNR1* | cannabinoid receptor 1 |
| *PTPRN2* | protein tyrosine phosphatase receptor type N2 |
| *SERP1* | stress associated endoplasmic reticulum protein 1 |
| *CD74* | CD74 molecule |
| *LRRC8A* | leucine rich repeat containing 8 VRAC subunit A |
| *PICK1* | protein interacting with PRKCA 1 |
| *EGFR* | epidermal growth factor receptor |
| *INS* | insulin |

| (b) Somatotropin: Pituitary gland (predictions of somatotropin-producing genes) | |
|---|---|
| Top genes | Gene names |
| *HDAC1* | histone deacetylase 1 |
| *EGFR* | epidermal growth factor receptor |
| *FKBP1B* | FKBP prolyl isomerase 1B |
| *FFAR4* | free fatty acid receptor 4 |
| *S100A8* | S100 calcium binding protein A8 |
| *PER2* | period circadian regulator 2 |
| *RFX3* | regulatory factor X3 |
| *PRKCE* | protein kinase C epsilon |
| *SERP1* | stress associated endoplasmic reticulum protein 1 |
| *ITSN1* | intersectin 1 |

Table S2: Gene names of the top 10 predicted genes by MultiCens (ranked only among genes involved in peptide secretion) for the two primary peptide hormones: (a) insulin, and (b) somatotropin. See also Fig. 4b in main text for more context.

| | Insulin | | | |
|---|---|---|---|---|
| | Pancreas | | Skeletal Muscle | |
| | lncRNA symbol | References | lncRNA symbol | References |
| 1 | LINC00672 | [10,11] | ZEB1-AS1 | [12,13,14,15] |
| 2 | HOXA-AS2 | [5,16,17] | TNK2-AS1 | [18] |
| 3 | PRR34-AS1 | [19,20] | PWAR6 | [19,21,22,23] |
| 4 | MIR22HG | [24] | PRRT3-AS1 | [25,26] |
| 5 | LINC00294 | [27] | PRKCQ-AS1 | [28,29] |

| | Somatotropin | | | |
|---|---|---|---|---|
| | Pituitary Gland | | Liver | |
| | lncRNA symbol | References | lncRNA symbol | References |
| 1 | LINC01588 | [30] | NEAT1 | [31] |
| 2 | PTPRD-AS1 | [32,33] | ZNF528-AS1 | [34] |
| 3 | LINC01132 | [35] | MIR210HG | [36,37] |
| 4 | UCA1 | [38] | ALMS1-IT1 | None |
| 5 | LINC01473 | [39] | LINC01278 | [40] |

| | Progesterone | | | |
|---|---|---|---|---|
| | Ovaries | | Uterus | |
| | lncRNA symbol | References | lncRNA symbol | References |
| 1 | CCDC18-AS1 | None | HAGLR | [41,42] |
| 2 | LINC00641 | [43,44,45] | TAF1A-AS1 | [46] |
| 3 | MIR210HG | [47,48,49,50] | LINC00602 | None |
| 4 | LINC01016 | [51,52] | PCAT19 | [53] |
| 5 | BEAN1-AS1 | None | HHIP-AS1 | [54] |

| | Norepinephrine | | | |
|---|---|---|---|---|
| | Adrenal Glands | | Small Intestine | |
| | lncRNA symbol | References | lncRNA symbol | References |
| 1 | PGM5P4-AS1 | None | RNF139-AS1 | None |
| 2 | CCDC18-AS1 | None | CARMN | [55] |
| 3 | MAGI2-AS3 | [56] | SPATA41 | [57] |
| 4 | LINC01291 | None | GHET1 | [58] |
| 5 | TOLLIP-AS1 | None | ATP1B3-AS1 | None |

Table S3: Top predicted lncRNAs for the hormones along with the references. These references show association of these lncRNAs to the corresponding hormone and related diseases.

### 4 Supplementary Figures

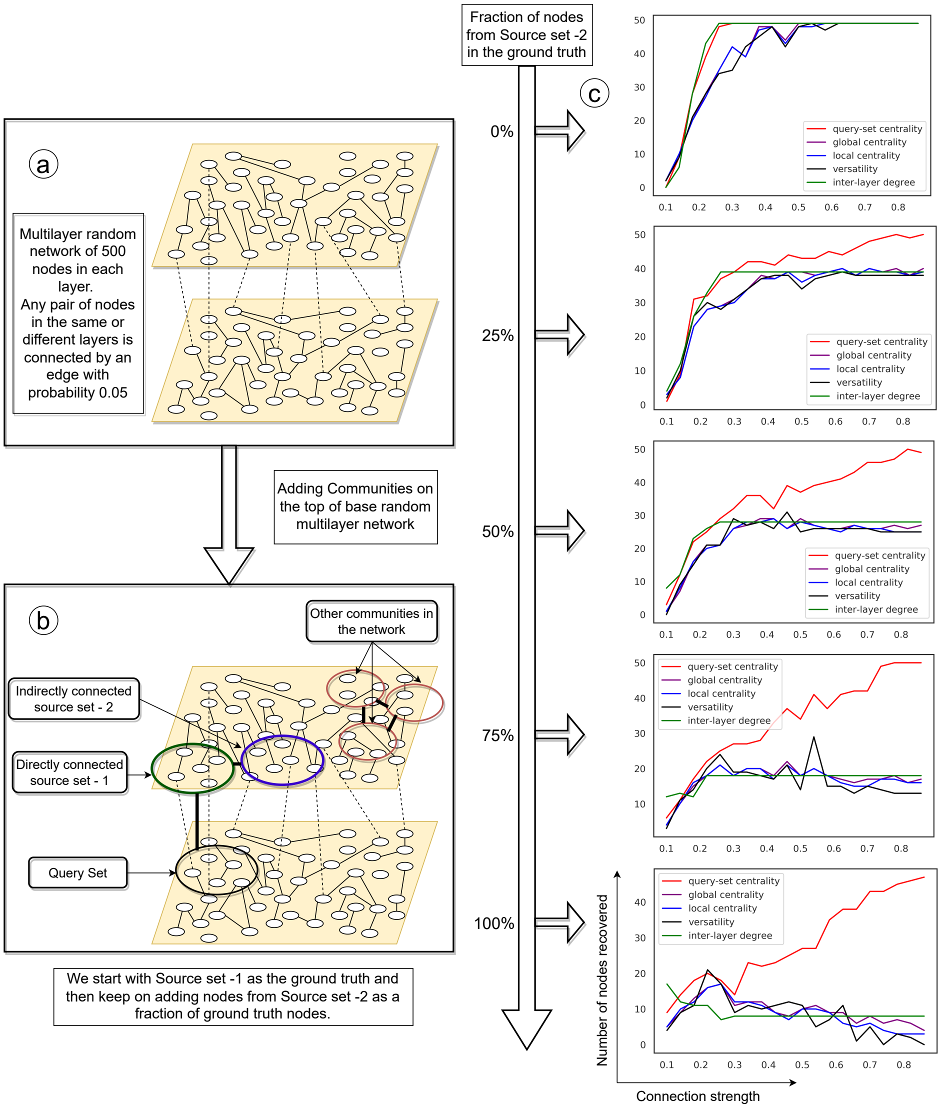

**Fig. S1.** (a,b) Synthetic network construction schematic, repeated from Fig. 2a,b of main text for convenience; (c) Results for local and global centrality to supplement the results shown in Fig. 2c of main text. As more nodes from *source-set-2* become part of the ground truth (shown as increasing percentages), our MultiCens query-set centrality outperforms the existing methods and MultiCens local and global centralities to a larger extent. Each plot shows the connection strength (x-axis) against the number of ground truth nodes in the top 100 ranked nodes (y-axis). Note that since the random graphs used here and in Fig. 2c of main text are not identical, the plots for the same measure between these figures are not identical too; however the performance trends are similar.

| Hormone Name | Number of Source Genes | Number of Target Genes | Retrieving Source genes | Retrieving Target genes |
|---|---|---|---|---|
| Aldosterone | 11 | 2 | 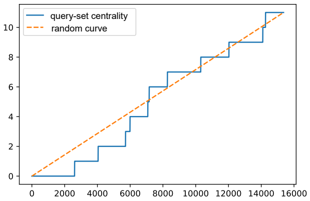  | 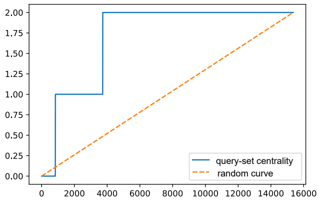  |
| Angiotensin-IV | 1 | 15 | 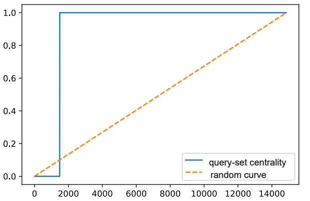  | 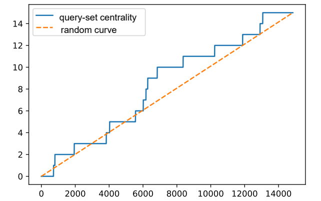  |
| Adrenaline | 4 | 24 | 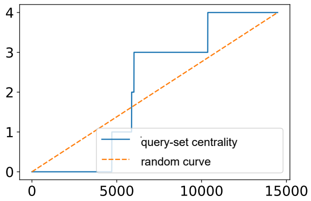  | 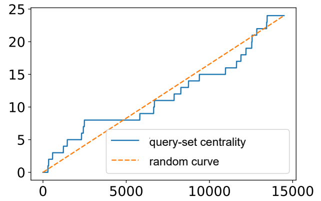  |
| Cortisol | 12 | 3 | 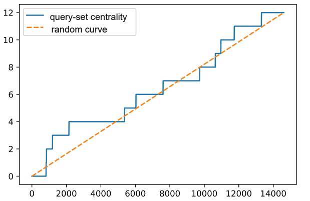  | 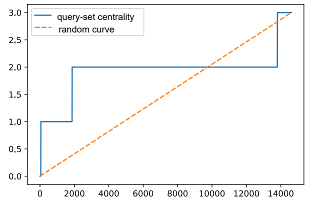  |
| Estradiol | 2 | 89 | 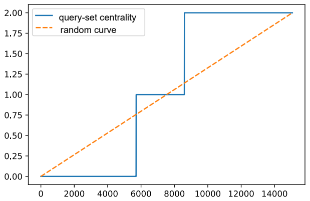  | 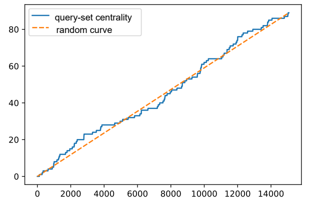  |
| Glucagon | 7 | 19 | 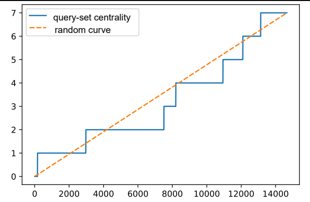  | 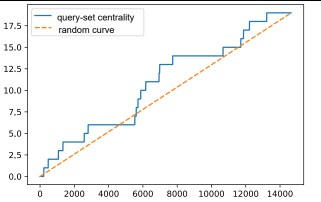  |

| Hormone Name | Number of Source Genes | Number of Target Genes | Retrieving Source genes | Retrieving Target genes |
|---|---|---|---|---|
| Thyroxin | 12 | 8 | 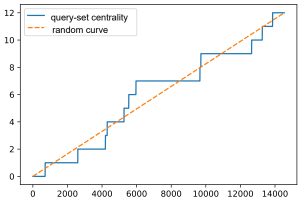 | 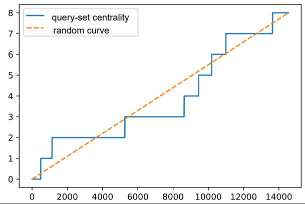 |
| Vitamin-D | 8 | 41 | 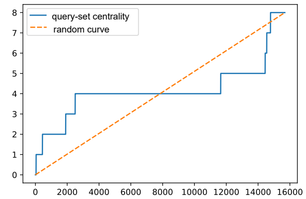 | 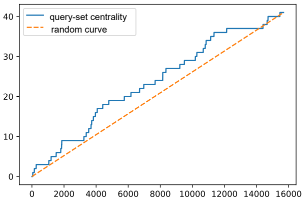 |

**Fig. S2.** Performance of MultiCens on all tested non-primary hormones, i.e., hormones with insufficient gene associations in the ground-truth in the following sense – these hormones have at least 10 genes in either the producing (source) set or the responding (target) set, but *not* in both sets unlike the primary hormones. See Fig. 3a of main text for similar recall-at-k curves for the primary hormones; Fig. 3b of main text and Suppl Table S1 summarizes the area under these curves for all tested hormones.

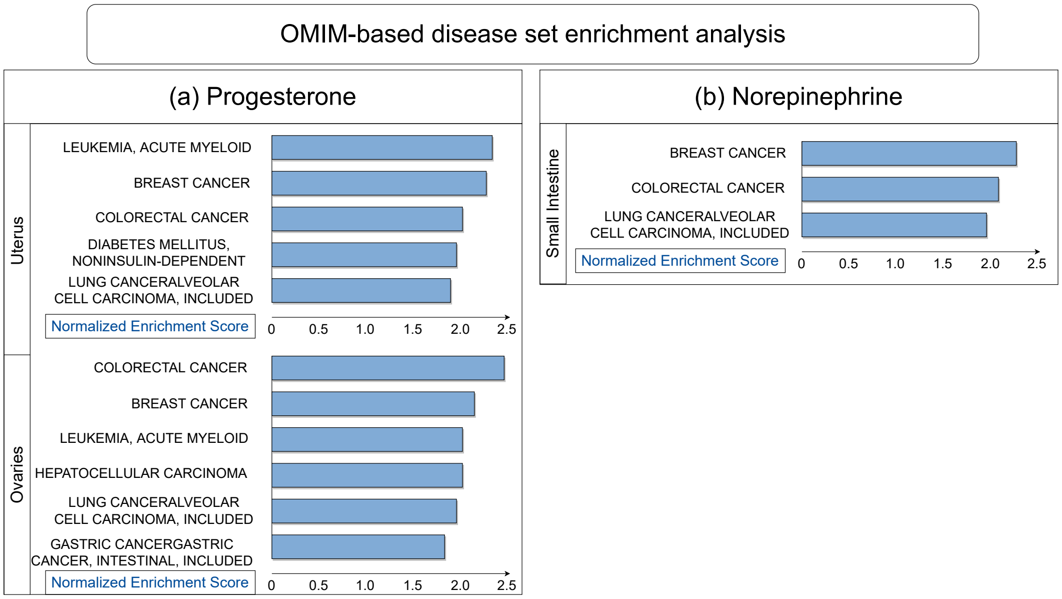

**Fig. S3.** OMIM-based disease set enrichment analysis of the centrality scores. We use WebGestalt to get these enrichments and apply an FDR cut-off of 0.05. For Norepinephrine, we do not see any significant enrichments at this FDR cutoff in Adrenal Glands. See also Fig. 4a in main text for similar enrichment analysis for the other two primary hormones: insulin and somatotropin.

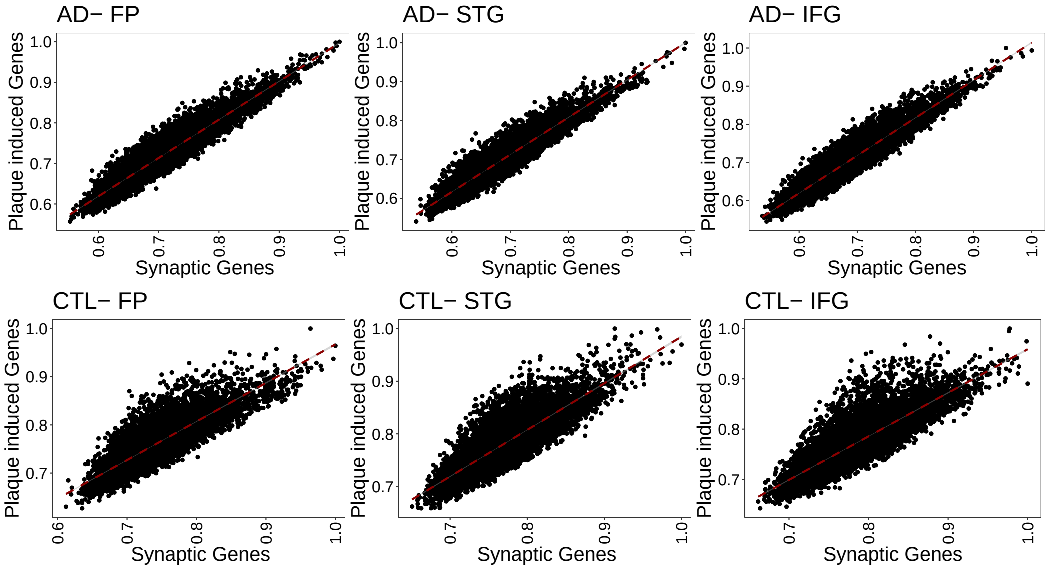

**Fig. S4.** Scatter plots representing correlation of centrality scores obtained using SSG vs PIG-based query sets. It can be inferred that the centrality scores with respect to different query sets shows less deviation in AD-based multilayer network than the control group.

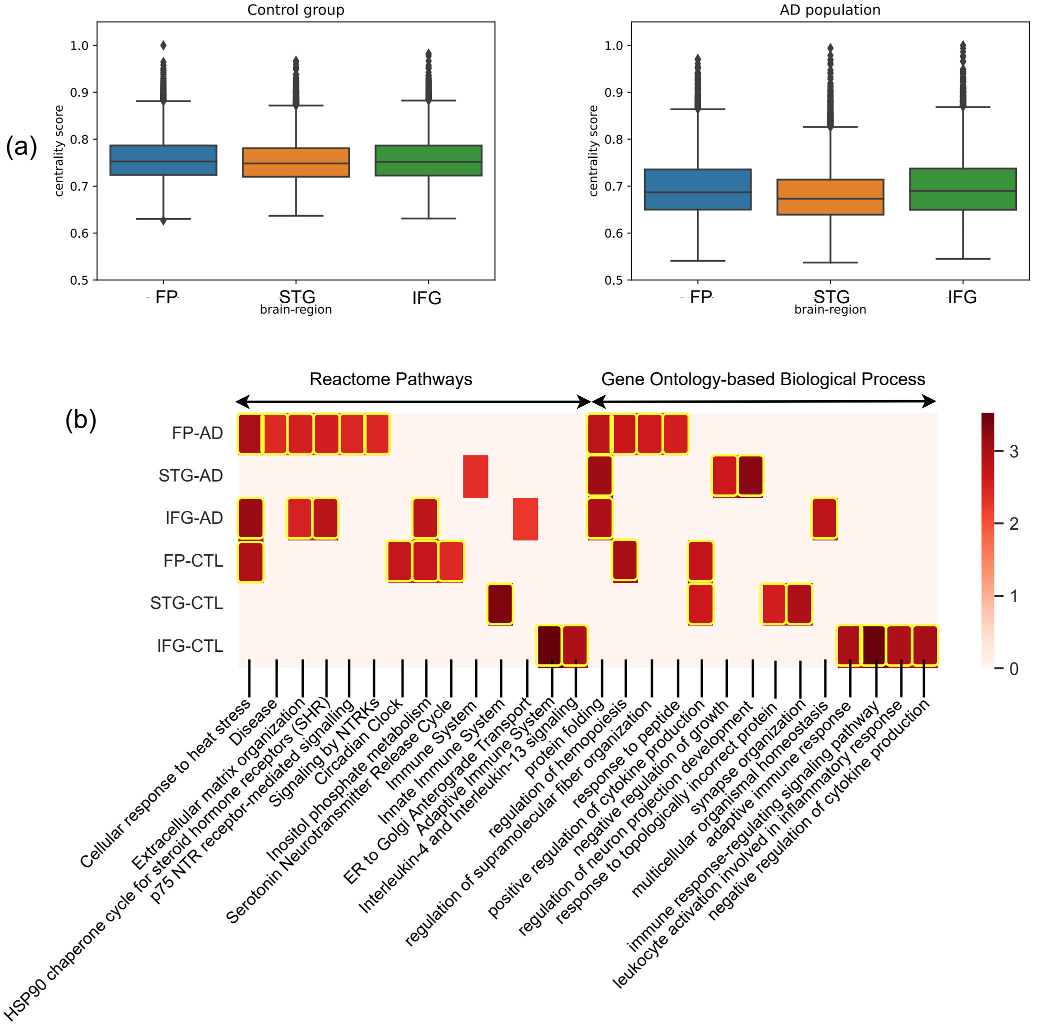

**Fig. S5. PIG-based query set**: Study of changes in the centrality-based gene rankings of four-layer networks of control and Alzheimer affected population. The PIG *query-set* is present in parahippocampal gyrus (PHG) and we rank genes of frontal pole (FP), superior temporal gyrus (STG) and inferior frontal gyrus (IFG). (a) Bar-plot showing region-wise shift of centrality scores of the three regions. (b) Reactome pathways and Gene Ontology-based process (GO-BP) enrichment analysis of each region in control and AD state. Color map represents the normalized enrichment score from WebGestalt. The highlighted boxes pass the 0.01 FDR cut-off. If centrality-based gene rankings of a region do not pass the 0.05 FDR cut off for an enrichment, we set the corresponding normalized enrichment score to 0.

#### 5 Supplementary Files

Suppl Dataset SD1: MultiCens on human multilayer networks, related to the four primary hormones. Link: https://github.com/BIRDSgroup/MultiCens/tree/main/results.

Suppl Dataset SD2: Results using brain region networks for AD and CTL populations. Link: https://github.com/BIRDSgroup/MultiCens/tree/main/brain_region_results.

#### References

1. L. Chung, A. E. Nelson, K. K. Ho, and R. C. Baxter, "Proteomic profiling of growth hormone-responsive proteins in human peripheral blood leukocytes," *J Clin Endocrinol Metab*, vol. 94, pp. 3038–3043, Aug 2009.
2. L. González, J. G. Miquet, P. E. Irene, M. E. Díaz, S. P. Rossi, A. I. Sotelo, M. B. Frungieri, C. M. Hill, A. Bartke, and D. Turyn, "Attenuation of epidermal growth factor (EGF) signaling by growth hormone (GH)," *J Endocrinol*, vol. 233, pp. 175–186, 05 2017.
3. E. Kostopoulou, A. P. Rojas-Gil, A. Karvela, and B. E. Spiliotis, "Epidermal growth factor receptor (EGFR) involvement in successful growth hormone (GH) signaling in GH transduction defect," *J Pediatr Endocrinol Metab*, vol. 30, pp. 221–230, Feb 2017.
4. D. Y. Oh and E. Walenta, "Omega-3 Fatty Acids and FFAR4," *Front Endocrinol (Lausanne)*, vol. 5, p. 115, 2014.
5. X. Li and H. M. Yu, "Overexpression of HOXA-AS2 inhibits inflammation and apoptosis in podocytes via sponging miRNA-302b-3p to upregulate TIMP3," *Eur Rev Med Pharmacol Sci*, vol. 24, pp. 4963–4970, 05 2020.
6. C. Dieter, N. E. Lemos, N. R. F. Corrêa, T. S. Assmann, and D. Crispim, "The Impact of lncRNAs in Diabetes Mellitus: A Systematic Review and In Silico Analyses," *Front Endocrinol (Lausanne)*, vol. 12, p. 602597, 2021.
7. Y. Sun, S. Xu, M. Jiang, X. Liu, L. Yang, Z. Bai, and Q. Yang, "Role of the Extracellular Matrix in Alzheimer's Disease," *Front Aging Neurosci*, vol. 13, p. 707466, 2021.
8. D. F. Weaver, "Amyloid beta is an early responder cytokine and immunopeptide of the innate immune system," *Alzheimers Dement (N Y)*, vol. 6, no. 1, p. e12100, 2020.
9. A. D. Sarma, A. R. Molla, G. Pandurangan, and E. Upfal, "Fast distributed pagerank computation," in *International Conference on Distributed Computing and Networking*, pp. 11–26, Springer, 2013.
10. W. Li, H. Li, L. Zhang, M. Hu, F. Li, J. Deng, M. An, S. Wu, R. Ma, J. Lu, *et al.*, "Long non-coding rna linc00672 contributes to p53 protein-mediated gene suppression and promotes endometrial cancer chemosensitivity," *Journal of Biological Chemistry*, vol. 292, no. 14, pp. 5801–5813, 2017.
11. D. Li, J. Lu, H. Li, S. Qi, and L. Yu, "Identification of a long noncoding rna signature to predict outcomes of glioblastoma," *Molecular medicine reports*, vol. 19, no. 6, pp. 5406–5416, 2019.
12. L. Gu, H. Sun, and Z. Yan, "lncrna zeb1-as1 is downregulated in diabetic lung and regulates lung cell apoptosis," *Experimental and Therapeutic Medicine*, vol. 20, no. 6, pp. 1–1, 2020.
13. Q. Meng, X. Zhai, Y. Yuan, Q. Ji, and P. Zhang, "lncrna zeb1-as1 inhibits high glucose-induced emt and fibrogenesis by regulating the mir-216a-5p/bmp7 axis in diabetic nephropathy," *Brazilian Journal of Medical and Biological Research*, vol. 53, no. 4, 2020.
14. Y. Song, C. Miao, and J. Wang, "Lncrna zeb1-as1 inhibits renal fibrosis in diabetic nephropathy by regulating the mir-217/mafb axis," *RSC advances*, vol. 9, no. 52, pp. 30389–30397, 2019.
15. G. Wei, T. Lu, J. Shen, and J. Wang, "LncRNA ZEB1-AS1 promotes pancreatic cancer progression by regulating miR-505-3p/TRIB2 axis," *Biochem Biophys Res Commun*, vol. 528, pp. 644–649, 08 2020.
16. Y. Lian, Z. Li, Y. Fan, Q. Huang, J. Chen, W. Liu, C. Xiao, and H. Xu, "The lncRNA-HOXA-AS2/EZH2/LSD1 oncogene complex promotes cell proliferation in pancreatic cancer," *Am J Transl Res*, vol. 9, no. 12, pp. 5496–5506, 2017.
17. F. X. Zheng, X. Q. Wang, W. X. Zheng, and J. Zhao, "Long noncoding RNA HOXA-AS2 promotes cell migration and invasion via upregulating IGF-2 in non-small cell lung cancer as an oncogene," *Eur Rev Med Pharmacol Sci*, vol. 23, pp. 4793–4799, Jun 2019.
18. L. Guo, H. Ma, Y. Kong, G. Leng, G. Liu, and Y. Zhang, "Long non-coding RNA TNK2 AS1/microRNA-125a-5p axis promotes tumor growth and modulated phosphatidylinositol 3 kinase/AKT pathway," *J Gastroenterol Hepatol*, vol. 37, pp. 124–133, Jan 2022.
19. G.-M. Liu, H.-D. Zeng, C.-Y. Zhang, and J.-W. Xu, "Key genes associated with diabetes mellitus and hepatocellular carcinoma," *Pathology-Research and Practice*, vol. 215, no. 11, p. 152510, 2019.

20. Z. Liu, Z. Li, B. Xu, H. Yao, S. Qi, and J. Tai, "Long Noncoding RNA PRR34-AS1 Aggravates the Progression of Hepatocellular Carcinoma by Adsorbing microRNA-498 and Thereby Upregulating FOXO3," *Cancer Manag Res*, vol. 12, pp. 10749–10762, 2020.
21. T. Nagai and M. Mori, "Prader-willi syndrome, diabetes mellitus and hypogonadism," *Biomedicine & pharmacotherapy*, vol. 53, no. 10, pp. 452–454, 1999.
22. R. Basheer, M. J. A. Jalal, and R. Gomez, "An unusual case of adolescent type 2 diabetes mellitus: Prader–willi syndrome," *Journal of family medicine and primary care*, vol. 5, no. 1, p. 181, 2016.
23. C. Zhang, H. Liu, P. Xu, Y. Tan, Y. Xu, L. Wang, B. Liu, Q. Chen, and D. Tian, "Identification and validation of a five-lncRNA prognostic signature related to Glioma using bioinformatics analysis," *BMC Cancer*, vol. 21, p. 251, Mar 2021.
24. Z. Lin, X. Li, X. Zhan, L. Sun, J. Gao, Y. Cao, and H. Qiu, "Construction of competitive endogenous rna network reveals regulatory role of long non-coding rnas in type 2 diabetes mellitus," *Journal of cellular and molecular medicine*, vol. 21, no. 12, pp. 3204–3213, 2017.
25. S. Yang, Y. Zhou, X. Zhang, L. Wang, J. Fu, X. Zhao, and L. Yang, "The prognostic value of an autophagy-related lncrna signature in hepatocellular carcinoma," *BMC bioinformatics*, vol. 22, no. 1, pp. 1–16, 2021.
26. L. Fan, H. Li, and W. Wang, "Long non-coding RNA PRRT3-AS1 silencing inhibits prostate cancer cell proliferation and promotes apoptosis and autophagy," *Exp Physiol*, vol. 105, pp. 793–808, 05 2020.
27. J. Qiu, S. Zhou, W. Cheng, and C. Luo, "LINC00294 induced by GRP78 promotes cervical cancer development by promoting cell cycle transition," *Oncol Lett*, vol. 20, p. 262, Nov 2020.
28. J. A. Timmons, P. J. Atherton, O. Larsson, S. Sood, I. O. Blokhin, R. J. Brogan, C.-H. Volmar, A. R. Josse, C. Slentz, C. Wahlestedt, *et al.*, "A coding and non-coding transcriptomic perspective on the genomics of human metabolic disease," *Nucleic acids research*, vol. 46, no. 15, pp. 7772–7792, 2018.
29. S. I. Itani, W. J. Pories, K. G. Macdonald, and G. L. Dohm, "Increased protein kinase C theta in skeletal muscle of diabetic patients," *Metabolism*, vol. 50, pp. 553–557, May 2001.
30. J. Volejnikova, P. Vojta, H. Urbankova, R. Mojzíkova, M. Horvathova, I. Hochova, J. Cermak, J. Blatny, M. Sukova, E. Bubanska, *et al.*, "Czech and slovak diamond-blackfan anemia (dba) registry update: Clinical data and novel causative genetic lesions," *Blood Cells, Molecules, and Diseases*, vol. 81, p. 102380, 2020.
31. S. S. Jin, C. J. Lin, X. F. Lin, J. Z. Zheng, and H. Q. Guan, "Silencing lncRNA NEAT1 reduces nonalcoholic fatty liver fat deposition by regulating the miR-139-5p/c-Jun/SREBP-1c pathway," *Ann Hepatol*, vol. 27, no. 2, p. 100584, 2022.
32. M. Zhou, Y. Sun, Y. Sun, W. Xu, Z. Zhang, H. Zhao, Z. Zhong, and J. Sun, "Comprehensive analysis of lncRNA expression profiles reveals a novel lncRNA signature to discriminate nonequivalent outcomes in patients with ovarian cancer," *Oncotarget*, vol. 7, pp. 32433–32448, May 2016.
33. G. R. Uhl and M. J. Martinez, "PTPRD: neurobiology, genetics, and initial pharmacology of a pleiotropic contributor to brain phenotypes," *Ann N Y Acad Sci*, vol. 1451, pp. 112–129, 09 2019.
34. E. Rothzerg, X. D. Ho, J. Xu, D. Wood, A. Märtson, and S. Kõks, "Upregulation of 15 antisense long non-coding rnas in osteosarcoma," *Genes*, vol. 12, no. 8, p. 1132, 2021.
35. W. Zhu, X. Xiao, and J. Chen, "Silencing of the long noncoding RNA LINC01132 alleviates the oncogenicity of epithelial ovarian cancer by regulating the microRNA-431-5p/SOX9 axis," *Int. J. Mol. Med.*, vol. 48, Aug. 2021.
36. Q. Sun, Y. J. Song, and K. V. Prasanth, "One locus with two roles: microrna-independent functions of microrna-host-gene locus-encoded long noncoding rnas," *Wiley Interdisciplinary Reviews: RNA*, vol. 12, no. 3, p. e1625, 2021.
37. Y. Wang, W. Li, X. Chen, Y. Li, P. Wen, and F. Xu, "MIR210HG predicts poor prognosis and functions as an oncogenic lncRNA in hepatocellular carcinoma," *Biomed Pharmacother*, vol. 111, pp. 1297–1301, Mar 2019.

38. G. Liu, L. Wang, and Y. Li, "Inhibition of lncRNA-UCA1 suppresses pituitary cancer cell growth and prolactin (PRL) secretion via attenuating glycolysis pathway," *In Vitro Cell. Dev. Biol. Anim.*, vol. 56, pp. 642–649, Sept. 2020.
39. F. Peng, S. Yan, H. Liu, Z. Liu, F. Jiang, P. Cao, and R. Fu, "Roles of LINC01473 and CD74 in osteoblasts in multiple myeloma bone disease," *J Investig Med*, Feb 2022.
40. W. J. Huang, X. P. Tian, S. X. Bi, S. R. Zhang, T. S. He, L. Y. Song, J. P. Yun, Z. G. Zhou, R. M. Yu, and M. Li, "The $\beta$-catenin/TCF-4-LINC01278-miR-1258-Smad2/3 axis promotes hepatocellular carcinoma metastasis," *Oncogene*, vol. 39, pp. 4538–4550, 06 2020.
41. L. Jin, C. Luo, X. Wu, M. Li, S. Wu, and Y. Feng, "Lncrna-haglr motivates triple negative breast cancer progression by regulation of wnt2 via sponging mir-335-3p," *Aging (Albany NY)*, vol. 13, no. 15, p. 19306, 2021.
42. J. Tang, J. Ren, Q. Cui, D. Zhang, D. Kong, X. Liao, M. Lu, Y. Gong, and G. Wu, "A prognostic 10-lncrna expression signature for predicting the risk of tumour recurrence in breast cancer patients," *Journal of cellular and molecular medicine*, vol. 23, no. 10, pp. 6775–6784, 2019.
43. N. Dastmalchi, R. Safaralizadeh, S. Latifi-Navid, S. M. Banan Khojasteh, B. Mahmud Hussen, and S. Teimourian, "An updated review of the role of lncrnas and their contribution in various molecular subtypes of breast cancer," *Expert Review of Molecular Diagnostics*, vol. 21, no. 10, pp. 1025–1036, 2021.
44. Q. Mao, M. Lv, L. Li, Y. Sun, S. Liu, Y. Shen, Z. Liu, and S. Luo, "Long intergenic noncoding RNA 00641 inhibits breast cancer cell proliferation, migration, and invasion by sponging miR-194-5p," *J Cell Physiol*, vol. 235, pp. 2668–2675, 03 2020.
45. X. Han and S. Zhang, "Role of Long Non-Coding RNA LINC00641 in Cancer," *Front Oncol*, vol. 11, p. 829137, 2021.
46. L. Jin, C. Li, T. Liu, and L. Wang, "A potential prognostic prediction model of colon adenocarcinoma with recurrence based on prognostic lncRNA signatures," *Hum Genomics*, vol. 14, p. 24, 06 2020.
47. X.-y. Li, L.-y. Zhou, H. Luo, Q. Zhu, L. Zuo, G.-y. Liu, C. Feng, J.-y. Zhao, Y.-y. Zhang, and X. Li, "The long noncoding rna mir210hg promotes tumor metastasis by acting as a cerna of mir-1226-3p to regulate mucin-1c expression in invasive breast cancer," *Aging (Albany NY)*, vol. 11, no. 15, p. 5646, 2019.
48. Y. Du, N. Wei, R. Ma, S.-H. Jiang, and D. Song, "Long noncoding rna mir210hg promotes the warburg effect and tumor growth by enhancing hif-1$\alpha$ translation in triple-negative breast cancer," *Frontiers in oncology*, vol. 10, 2020.
49. W. Shi, Y. Tang, J. Lu, Y. Zhuang, and J. Wang, "MIR210HG promotes breast cancer progression by IGF2BP1 mediated m6A modification," *Cell Biosci*, vol. 12, p. 38, Mar 2022.
50. J. Ma, F. F. Kong, D. Yang, H. Yang, C. Wang, R. Cong, and X. X. Ma, "lncRNA MIR210HG promotes the progression of endometrial cancer by sponging miR-337-3p/137 via the HMGA2-TGF-$\beta$/Wnt pathway," *Mol Ther Nucleic Acids*, vol. 24, pp. 905–922, Jun 2021.
51. X. Li, F. Jin, and Y. Li, "A novel autophagy-related lncrna prognostic risk model for breast cancer," *Journal of Cellular and Molecular Medicine*, vol. 25, no. 1, pp. 4–14, 2021.
52. X. Pan, D. Li, J. Huo, F. Kong, H. Yang, and X. Ma, "Linc01016 promotes the malignant phenotype of endometrial cancer cells by regulating the mir-302a-3p/mir-3130-3p/nfya/satb1 axis," *Cell death & disease*, vol. 9, no. 3, pp. 1–18, 2018.
53. J. T. Hua, M. Ahmed, H. Guo, Y. Zhang, S. Chen, F. Soares, J. Lu, S. Zhou, M. Wang, H. Li, N. B. Larson, S. K. McDonnell, P. S. Patel, Y. Liang, C. Q. Yao, T. van der Kwast, M. Lupien, F. Y. Feng, A. Zoubeidi, M. S. Tsao, S. N. Thibodeau, P. C. Boutros, and H. H. He, "Risk SNP-Mediated Promoter-Enhancer Switching Drives Prostate Cancer through lncRNA PCAT19," *Cell*, vol. 174, pp. 564–575, 07 2018.
54. N. Li and X. Zhan, "Identification of clinical trait–related lncrna and mrna biomarkers with weighted gene co-expression network analysis as useful tool for personalized medicine in ovarian cancer," *EPMA Journal*, vol. 10, no. 3, pp. 273–290, 2019.

55. K. Dong, J. Shen, X. He, G. Hu, L. Wang, I. Osman, K. M. Bunting, R. Dixon-Melvin, Z. Zheng, H. Xin, M. Xiang, A. Vazdarjanova, D. J. R. Fulton, and J. Zhou, "Is an Evolutionarily Conserved Smooth Muscle Cell-Specific LncRNA That Maintains Contractile Phenotype by Binding Myocardin," *Circulation*, vol. 144, pp. 1856–1875, 12 2021.
56. S. Ghosal, B. Zhu, T.-T. Huynh, L. Meuter, A. Jha, S. Talvacchio, M. Knue, M. Patel, T. Prodanov, S. Das, *et al.*, "A long noncoding rna–microrna expression signature predicts metastatic signature in pheochromocytomas and paragangliomas," *Endocrine*, pp. 1–10, 2021.
57. Y. Zhang, X. You, S. Li, Q. Long, Y. Zhu, Z. Teng, and Y. Zeng, "Peripheral blood leukocyte rna-seq identifies a set of genes related to abnormal psychomotor behavior characteristics in patients with schizophrenia," *Medical science monitor: international medical journal of experimental and clinical research*, vol. 26, pp. e922426–1, 2020.
58. S. Li, J. Liu, F. Kong, Y. Wang, N. Li, and Y. Zou, "lncRNA GHET1 has effects in development of pre-eclampsia," *J Cell Biochem*, vol. 120, pp. 12647–12652, 08 2019.
